# Sleep and Activity Patterns in Autism Spectrum Disorder

**DOI:** 10.1101/2024.05.02.592263

**Authors:** J. Dylan Weissenkampen, Arpita Ghorai, Maria Fasolino, Brielle Gehringer, Maya Rajan, Holly C. Dow, Ronnie Sebro, Daniel J. Rader, Brendan T. Keenan, Laura Almasy, Edward S. Brodkin, Maja Bucan

**Affiliations:** Department of Genetics, University of Pennsylvania; Department of Psychiatry, University of Pennsylvania; Department of Radiology, Mayo Clinic; Division of Sleep Medicine, University of Pennsylvania; Department of Medicine, University of Pennsylvania; Department of Biomedical and Health Informatics, Children’s Hospital of Philadelphia

**Author notes:** Corresponding author: Maja Bucan.

**Keywords:** Autism Spectrum Disorder, actimetry, sleep, physical activity, circadian

## Abstract

**Background:** Autism spectrum disorder (ASD) is a highly heritable and heterogeneous neurodevelopmental disorder characterized by impaired social interactions, repetitive behaviors, and a wide range of comorbidities. Between 44-83% of autistic individuals report sleep disturbances, which may share an underlying neurodevelopmental basis with ASD.

**Methods:** We recruited 382 ASD individuals and 223 of their family members to obtain quantitative ASD-related traits and wearable device-based accelerometer data spanning three consecutive weeks. An unbiased approach identifying traits associated with ASD was achieved by applying the elastic net machine learning algorithm with five-fold cross-validation on 6,878 days of data. The relationship between sleep and physical activity traits was examined through linear mixed-effects regressions using each night of data.

**Results:** This analysis yielded 59 out of 242 actimetry measures associated with ASD status in the training set, which were validated in a test set (AUC: 0.777). For several of these traits (e.g. total light physical activity), the day-to-day variability, in addition to the mean, was associated with ASD. Individuals with ASD were found to have a stronger correlation between physical activity and sleep, where less physical activity decreased their sleep more significantly than that of their non-ASD relatives.

**Conclusions:** The average duration of sleep/physical activity and the variation in the average duration of sleep/physical activity strongly predict ASD status. Physical activity measures were correlated with sleep quality, traits, and regularity, with ASD individuals having stronger correlations. Interventional studies are warranted to investigate whether improvements in both sleep and increased physical activity may improve the core symptoms of ASD.

## INTRODUCTION

Autism spectrum disorder (ASD) is a childhood-onset neurodevelopmental disorder characterized by repetitive and stereotyped behaviors as well as deficits in social communication^1^. Along with these core defining symptoms of ASD, individuals frequently present with a plethora of other symptoms with varying severity, such as intellectual disability (ID), epilepsy, sensory deficits, gastrointestinal problems, motor difficulties, and sleep disturbances^2–4^.

Numerous small-scale studies and meta-analyses suggest that sleep disturbances are common in individuals with ASD^5^. Parental reports indicate a higher prevalence rate of disturbed sleep in children with autism (44-83%)^6,7^ compared to age-matched controls (25-40%)^8,9^. A questionnaire-based study of children with Asperger syndrome demonstrated an almost two-fold increase in the prevalence of short sleep duration and a more than five-fold higher prevalence of sleep onset problems compared to neurotypical age-matched controls^10^. Sleep problems were also among the most frequently reported clinical features (39.4%) in a recent study examining lifetime medical and psychiatric co-morbidities in 2,917 autistic subjects within the Simons Powering Autism Research (SPARK) cohort^11^. Autistic individuals have difficulties with a broad range of sleep traits, including bedtime resistance, sleep onset delay, night awakenings, parasomnias, sleep-disordered breathing, daytime sleepiness, and reduced “restorative value of sleep” ^12^. Furthermore, polysomnography studies have reported shorter total sleep time, longer sleep onset latency, and lower sleep efficiency in children with ASD ^7,12^. One research group recently showed that increased core autism spectrum traits and executive function impairment were associated with disruption of several sleep-wake parameters in adults, particularly daily sleep-wake rhythm. Moreover, increased executive function impairment was associated with disrupted sleep quality and level of physical activity^13^. This supports initial reports on the association between ASD symptom severity and quality of sleep ^13,14^.

Similar to sleep, motor abnormalities may often be an early developmental sign of ASD^15–17^. Motor disturbances observed in ASD are broad, encompassing the diagnostic “restrictive, repetitive stereotyped behaviors” as well as impaired fine motor control, coordination, and gait^18^. Wrist-worn accelerometers are increasingly being used for the assessment of physical activity and to derive indicators of sleep behavior, including, but not limited to, sleep duration, daytime inactivity, sleep efficiency, and number of sleep bouts^19^. In studies of non-clinical populations, individuals exhibiting lower levels of daytime activity have poorer sleep quality, while those with moderate-to-high levels of daytime physical activity have fewer sleep problems as measured by the Pittsburgh Sleep Quality Index (PSQI)^20–22^. This association between activity and sleep quality has also been observed in ASD, though the directionality of this relationship is unclear based on data from previous studies^23,24^. However, research focusing on activity levels in ASD has predominately centered on children, revealing decreased activity levels^25^. Notably, interventions aimed at increasing physical activity among individuals with ASD have shown promise in reducing deficits in social functioning tasks^26^. This provides evidence that exploring the sleep and physical activity patterns in ASD may provide insights into areas for focused and personalized interventions for improving core, defining ASD symptomology.

The data currently available on sleep in children with ASD are restricted by several experimental limitations that also complicate interpretation. First, many studies are based on subjective data, gathered from parental questionnaires, rather than objective measures of sleep^27^. Second, the number of children evaluated in any given study using actimetry or other objective measures has been small, with frequently less than one hundred per study ^27,28^. Third, because of conflicting data, it is unclear whether children with ASD and ID differ from those with ASD without ID with respect to the degree of sleep disturbance ^7,29–32^. Core ASD behavioral phenotypes and sleep/circadian disruption may have common underlying mechanisms, such as disrupted neuronal connectivity ^33^. During sleep, neuronal synapses undergo widespread alterations. During sleep, neuronal synapses undergo widespread alterations in composition and signaling ^34–36^, and sleep and wake cycles, along with physical activity ^37^ play a key role in regulating brain plasticity ^38,39^. Although the role of sleep in brain development is not fully understood, sleep promotes synapse formation, stabilization, and pruning during development ^40–43^. Therefore, sleep may be an important phenotypic assay for dissecting neuronal connectivity and analyzing the biological heterogeneity of ASD. We hypothesize that sleep and physical activity, and their interplay will be different in individuals with ASD than non-ASD family members.

To evaluate sleep, circadian behavior, and physical activity in ASD, we collected accelerometer-based data and ASD-related phenotypic data ^44,45^ on individuals with ASD and their families. With the use of machine learning methods and linear mixed effects regressions, we established relationships between sleep and physical activity trait disturbances and core ASD traits in the Autism Spectrum Program of Excellence (ASPE)^44^ and SPARK collections.

## METHODS & MATERIALS

### Study Participants

We recruited participants through the Autism Spectrum Program of Excellence (ASPE) at the University of Pennsylvania in Philadelphia, Pennsylvania ^13,44,45^, and 190 ASD participants from the Simons Powering Autism Research (SPARK) collection^46^. As our study focused on the analysis of ASD without intellectual disability (ID), we excluded subjects with ID (full-scale IQ less than 70, as estimated by the Shipley-2 ^47^). While comorbid psychiatric diagnoses per se were not an exclusion criterion, the following were exclusion criteria: 1) history of intellectual disability (ID), 2) recent (last four weeks) severe mood or psychotic symptoms, 3) recent severe aggressive or self-injurious behaviors, and 4) history of major neurological disorder (e.g.: dementia, severe head trauma, recent seizures). Each ASPE participant completed a 1-hour phone interview to collect general demographic information (see Supplementary Table 1) and assess psychiatric and developmental history, social behavior and autism spectrum traits to determine if they met criteria for autism spectrum disorder (ASD) as defined in the Diagnostic and Statistical Manual of Mental Disorders, Fifth Edition (DSM-5)^1^. This assessment identified probands and their family members who met the criteria for ASD according to a DSM-5 diagnostic checklist or other psychiatric disorders, as well as family members who did not meet criteria for such disorders ^44^. All probands and family members were also assessed using informant-reporting (most often parent) for the Social Responsiveness Scale (SRS) ^48,49^, the Broad Autism Phenotype Questionnaire (BAPQ) ^50^, and the Behavior Rating Inventory of Executive Function (BRIEF) ^51^. For those who passed these criteria (n = 522), we collected three weeks of actimetry data. Participants wore a GENEActiv accelerometer (Activinsights, Cambridgeshire, United Kingdom) for 21 days. The open-source R-package GGIR (version 1.10-7) was used to derive day-to-day sleep and activity data^52^. In the final analysis, we included data from participants with a minimum 5 days of recording data and at least 20 hours of actimetry data per day. Days were excluded from the analysis based on standard GGIR quality control^52^ and participants with fewer than three hours of sleep or who consistently on average went to sleep before 8 pm were also excluded. The within-individual, across-days mean and standard deviation of 109 actimetry-derived traits were obtained from GGIR, yielding 218 total variables. We analyzed the same filtered dataset with Accelerometer, a program specifically developed to obtain refined physical activity measures ^53–55^. The means and standard deviations for 12 Accelerometer-derived traits were included in our analysis, yielding an additional 24 variables. All actimetry-derived traits from GGIR and Accelerometer were then converted to z-scores using the whole sample as the distribution. In addition to the actimetry, participants kept a sleep diary that recorded their quality of sleep, time they went to bed, and time they awoke each day.

Further QC was performed based on the actimetry results in ASPE: seven participants were removed due to low wear time; 24 were lost to an error in the watches; 9 were removed for sleeping less than three hours a day; 17 were removed due to fewer than five days of data; 16 were lost due to bad data, nonwear, or night worker status; and 22 were removed due to improper consent, missing demographics data, or unconfirmed ASD status. Therefore, 415 (79.5%) of the participants were included in the final analyses, where 192 ASPE were ASD participants and 223 were family members who did not have ASD. This study was approved by the Institutional Review Board of the University of Pennsylvania.

In addition to the ASPE participants, we performed accelerometer-based evaluation on 201 ASD participants recruited through the SPARK Research Match The SPARK ^46^. This collection includes ASD status for all participants (n = 17,342 individuals from 4,770 families) and limited phenotype questionnaire-based data. 11 participants were excluded from the analysis based on: 5 daytime sleepers, 1 who did not sleep, 1 who was caring for an infant, 2 night workers, and 2 participants with watch issues, leading to 190 total SPARK ASD participants included. SPARK participants were further separated by whether they had a reported sleep problem, determined by if they reported a diagnosis of a sleep disorder or an undiagnosed sleep problem in the medical questionnaire. Overall, 142 SPARK ASD individuals reported no sleep problem and 48 reported a sleep problem. All ASPE and SPARK ASD subjects wore GENEActiv devices for 3 weeks and completed assessment tools listed above (SRS, BRIEF and BAPQ), in addition to a commonly used sleep assessment tool: the PROMIS Sleep Disturbance Measure–Short Form, a brief eight-item self-report measure that assesses sleep quality and disturbances over the last week ^56,57^.

### Machine Learning Using Elastic Net

To assess the potential association between actimetry-derived sleep and physical activity traits with ASD status, we employed the elastic net method of machine learning ^58,59^ on all GGIR and Accelerometer-derived traits. By systematically adjusting a penalization term, the elastic net logistic regression algorithm enabled us to explore a wide range of regression models and effectively eliminate irrelevant variables, reducing them to zero. To discern how to vary the loss function for the elastic net, α, 1,000 iterations of the machine learning were performed at values of α ranging from 0-1 in steps of 0.05. The best-performing α cutoff was determined as the cutoff with the highest median AUC (Supplementary Figure 2).

We employed complementary approaches for model validation. First, days with actimetry data were sequentially numbered (starting at day 1 for first day of recording) and split into two datasets – even and odd days. Data from odd days were used for training and validation of the model, utilizing five-fold cross-validation approach. Briefly, participants were randomly split into five groups, and the model was trained on the data from four of the groups and then validated on the data from the fifth group. Data from even days served as a separate quasi-independent testing set to control for potential overfitting of the models. The mean and standard deviation for each actimetry-derived sleep and physical activity trait were calculated separately within the odd day and even day datasets to evaluate whether the overall average and intra-individual variance of a trait differed in the two datasets. As ASD was a binary outcome variable, model error was determined and compared using binomial deviance.

### Association between Physical Activity and Sleep Traits

Next, we investigated the relationship between physical activity and sleep traits and evaluated whether ASD status modifies these relationships. We used a linear mixed effects regression for all ten total sleep traits derived from the GGIR and Accelerometer algorithms. Each 24-hour day/night cycle for every participant was included, with quality control filtering identical to the previous machine learning step. A linear mixed effects model was created for each trait combination using the lme4 R package ^60^, with the following covariates: age, age^2^, sex, ASD status, the physical activity trait of interest, the interaction between ASD and the physical activity trait, and two random effect variables to account for repeated measures within individuals (e.g., multiple days) and potential family effects. The p-values were derived from the t-scores in these models.

### A comparison of sleep and activity in ASPE and SPARK probands and family members

To further validate the machine learning findings in a separate population, we performed traditional regression analyses on 218 GGIR accelerometer-derived sleep and activity characteristics in 192 ASPE ASD probands (AP), 54 relatives of ASPE probands with a different psychiatric condition (AR), 80 relatives of ASD probands without a psychiatric diagnosis (UR), 142 SPARK subjects with ASD and no reported sleep problems (SP (noSP)), and 48 SPARK subjects with ASD who did report sleep problems (SP (wSP)). Participants recruited through *NRXN1* status were excluded from this analysis (n = 89). Individual linear regressions were performed for each trait as an outcome variable, with age, age^2^, and sex as covariates, and status as the predictor of interest. The status comparisons were done in a pair-wise manner. Residuals of the actimetry-derived sleep traits after regressing out age, age^2^, and sex were calculated and graphed as violin plots.

## RESULTS

### Analysis Sample and Actigraphy Data

192 ASD subjects and 223 non-ASD relatives (first and second-degree) in ASPE and 190 ASD subjects were included in the final analysis (demographics in Supplementary Table I). Overall research strategy is seen in Fig. 1a. On average, we analyzed 17.88 days per participant (median 17 days), with a mean (SD) of 23.6 (0.78) hours of data per day after excluding those with fewer than 5 days of data (red line) (Fig. 1b). The sleep duration during the nights was normally distributed with a mean of 6.80 hours (median of 6.82 hours) and a standard deviation of 1.67 hours (Fig. 1c). For the nights, the majority of nights for individuals had a median sleep efficiencies over 86.8% (Fig. 1d). We also collected sleep diaries for each subject. Among similar traits derived from the GENEActiv and GENEA data In R (GGIR) analysis (actimetry-derived data)^52^ and from the sleep diary (self-reported data), total sleep time, sleep offset, and sleep onset were moderately to highly correlated (r=0.44, 0.68, 0.73; p=XX). In contrast, sleep efficiency was poorly correlated between self-reported and actimetry-derived data (r=0.038).

**Figure 1.**
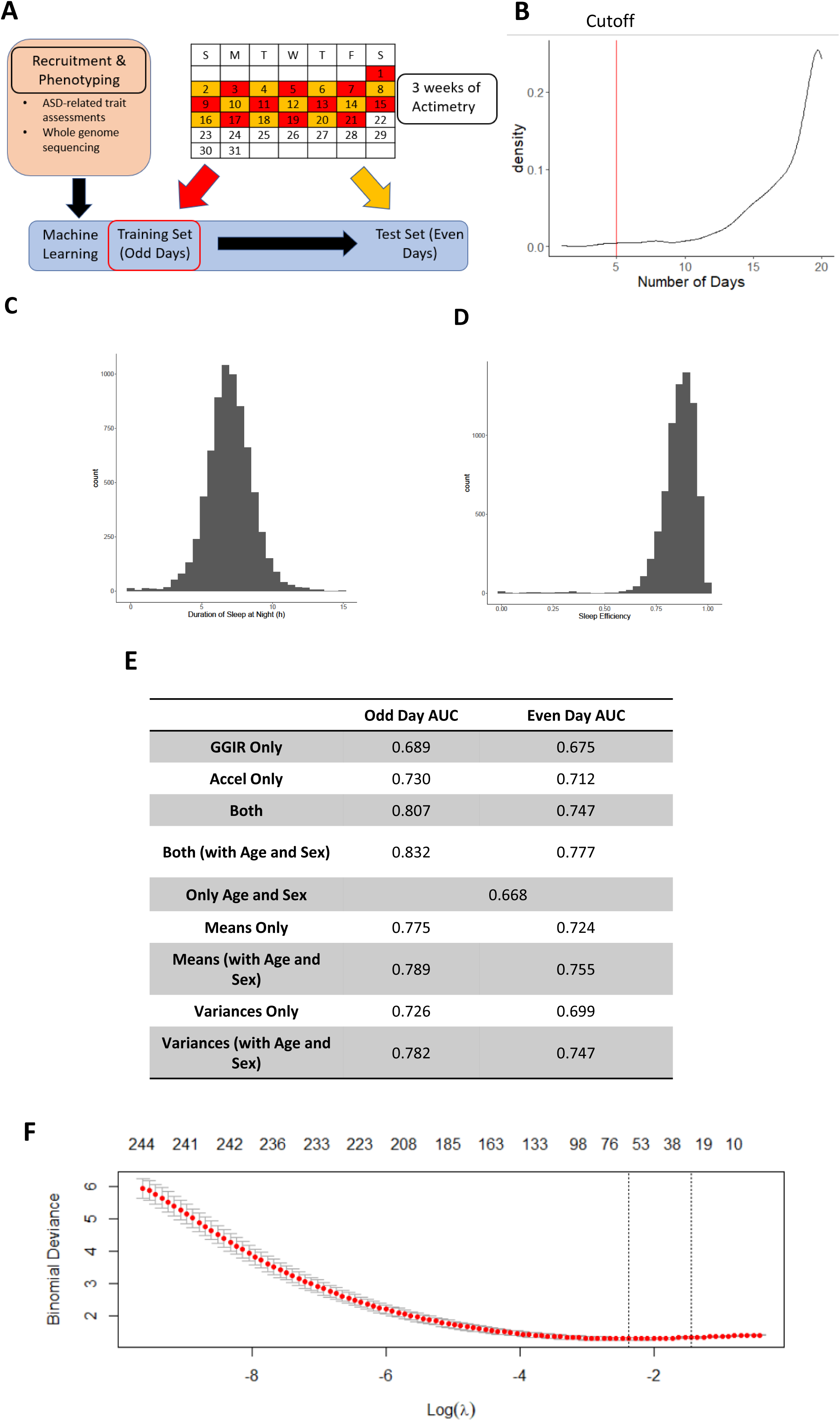
Prioritization of accelerometer-derived sleep and activity traits using machine learning. A) ASPE study subjects were recruited; their ASD-associated behavioral traits were assessed using questionnaires and their sleep and activity patterns were derived from 3 weeks of accelerometer-based data. The accelerometer-based data were split into two sets; even and odd days, where the odd days were used for elastic net to build a prediction model for ASD. The best-performing model was then evaluated in the even days dataset. B) The density plot for the number of days each participant has actimetry data which passes quality control. The cutoff (in red) shows 5 days. C) Distribution of total nighttime sleep for all participants. D) Total sleep efficiency distribution for all participants. E) The areas under the curve (AUCs) in the best-performing models for even and odd days. A significant difference was found between the demographics only model and the model including demographics and actimetry data, showing a significant improvement of prediction when actimetry is included (DeLong’s test, p< 0.01). There were not significant differences between the means only and variances only models (lowest p > 0.1). F) Plot showing model selection in elastic net using 5-fold cross-validation. Models were evaluated using binomial deviance. The top numbers refer to the number of variables within the model.

### Prioritization of sleep and physical activity traits using machine learning

To understand which actimetry-derived traits were associated with ASD, we utilized a machine learning approach on 242 actimetry-derived sleep and physical activity traits derived from the means and the day-to-day variations in the 171 actimetry-derived traits, age, and sex (Supplementary Table II; correlations in Supplementary Fig. 1). Through elastic net, we constructed models to predict ASD status by discerning which variables are most strongly associated with ASD. To evaluate what is the best hyperparameter value, α, we ran 1,000 iterations of the elastic net for each α value between 0 and 1, increasing by 0.05 (Supplementary Fig. 2). This value corresponds to the fraction the elastic net follows the ridge regression versus the least absolute shrinkage and selection operator (LASSO) for the loss function. The α value with the highest median AUC was selected. For the model including age and sex, 0.2 was selected for α, while 0.3 was selected for the models excluding age and sex. To control for overfitting, we employed two strategies: a 5-fold cross-validation for independent testing for model selection, and a quasi-independent sample by splitting the data into even and odd days and testing on the even days (Fig. 1a). There was a similar distribution of weekdays and weekend days for subjects with ASD and those without ASD. However, there were slight differences in the seasonality of the days collected for subjects with ASD and those without ASD, where individuals with ASD more commonly were measured in spring (ASD: 23.7%, no ASD: 18.8%; p < 0.001) and less commonly in summer (ASD: 21.3%, no ASD: 30.0%, p < 0.001) (Supplementary Fig. 3 & 4). The best-fitting model had areas under the curve (AUCs) of 0.832 and 0.777 in the odd and even day datasets, respectively (Fig. 1e,f; Table I). We performed a similar analysis without including age and sex in the models to evaluate how the actimetry traits perform alone. The best model had AUC of 0.807 and 0.747, in odd and even datasets, respectively (Supplementary Table III). Additionally, we used the odd day dataset to evaluate only demographic data (age and sex) as predictors in elastic net, as the demographic data did not change between even and odd days, even days could not be used as a validation set. This model gave an AUC of 0.668, which was significantly lower compared to the full model with actimetry traits included (AUC: 0.832; lower (DeLong’s test, p < 1e^-8^). This suggests that although age and sex were strong predictors of ASD status in our sample, the actimetry-derived traits significantly improved the model. The magnitudes of the effect sizes or beta values are proportional to the importance of the actimetry-derived trait in predicting ASD, as they were scaled to the z-scores by subtracting by the mean of the trait, divided by the standard deviation of the trait.

**Table I.** Elastic Net-Selected Actimetry-Derived Traits.

| Beta | Name |
| --- | --- |
| -0.1836 | Total vigorous activity (Mean) |
| -0.12399 | Total wear time (Mean) |
| -0.09339 | Average acceleration during unbouted inactivity (Mean) |
| -0.09039 | Time spent sedentary (SD) |
| -0.06819 | Time in vehicles (SD) |
| -0.06782 | Average acceleration during inactivity (SD) |
| -0.059 | Sleep duration in the sleep period (Mean) |
| -0.05575 | Time in mixed physical activity (SD) |
| -0.04805 | Duration of unbouted light activity (Mean) |
| -0.04663 | Total light physical activity (Mean) |
| -0.04504 | Number of blocks of unbouted inactivity (Mean) |
| -0.04461 | Number of blocks of unbouted light activity (SD) |
| -0.04252 | Number of blocks of unbouted inactivity (SD) |
| -0.03611 | Duration of unbouted light activity (SD) |
| -0.03578 | Duration of the day and sleep period (Mean) |
| -0.02982 | Number of blocks of unbouted light activity (Mean) |
| -0.02736 | Time in mixed physical activity (Mean) |
| -0.01703 | Age |
| -0.0141 | Duration of unbouted inactivity (SD) |
| -0.00899 | Average light during M10 (SD) |
| -0.00842 | Average light during M10 (Mean) |
| -0.00322 | Total vigorous activity (SD) |
| 0.003198 | Average acceleration during unbouted vigorous activity (SD) |
| 0.004062 | Average acceleration during moderate activity during the sleep period (SD) |
| 0.004919 | Duration of light activity during sleep period (Mean) |
| 0.006017 | Average acceleration during inactivity during the sleep period (Mean) |
| 0.007454 | Number of wakes of at least 5 minutes (Mean) |
| 0.007483 | Accumulated sustained inactivity bouts during the day (SD) |
| 0.009516 | Number of bouts of inactivity of 10-20 minutes (Mean) |
| 0.011149 | Number of blocks of inactivity in bouts of 10-20 minutes (Mean) |
| 0.011656 | Guider used wakeup time (SD) |
| 0.015455 | Number of bouts of light activity of at least 10 minutes (Mean) |
| 0.015484 | Number of blocks of light activity in bouts of at least 10 minutes (Mean) |
| 0.017182 | Sleep duration in the sleep period (SD) |
| 0.018188 | Number of blocks of moderate or vigorous activity in bouts of at least 10 minutes (Mean) |
| 0.018805 | Number of sustained inactivity bouts during the day (Mean) |
| 0.019053 | Duration of light activity bouts of at least 10 minutes (SD) |
| 0.020203 | Duration of moderate or vigorous activity bouts of 5-10 minutes (Mean) |
| 0.02074 | Average acceleration during light activity (SD) |
| 0.022713 | Average acceleration during inactivity in bouts of at least 30 minutes (Mean) |
| 0.02451 | Number of bouts of moderate or vigorous activity of at least 10 minutes (Mean) |
| 0.030561 | Number of blocks of moderate or vigorous activity in bouts of 5-10 minutes (Mean) |
| 0.031406 | Duration of light activity bouts of at least 10 minutes (Mean) |
| **0.032764** | Accumulated sustained inactivity bouts during the day (Mean) |
| **0.03695** | Maximum light during M5 (SD) |
| **0.036984** | Number of bouts of moderate or vigorous activity of 5-10 minutes (Mean) |
| **0.038387** | Number of bouts of inactivity of at least 30 minutes (Mean) |
| **0.038875** | Average acceleration during vigorous activity (SD) |
| **0.03922** | Number of blocks of inactivity in bouts of at least 30 minutes (Mean) |
| **0.041014** | Time spent sedentary (Mean) |
| **0.042543** | Duration of moderate or vigorous activity bouts of at least 10 minutes (Mean) |
| **0.046363** | Average acceleration during inactivity in bouts of 10-20 minutes (SD) |
| **0.051135** | M10 onset time(Mean) |
| **0.058658** | Guider used sleep period duration (SD) |
| **0.06179** | Duration of the sleep period (SD) |
| **0.064039** | M5 onset time (Mean) |
| **0.065005** | L10 onset time (SD) |
| **0.06769** | Average acceleration during moderate or vigorous activity in bouts of 1-5 minutes (SD) |
| **0.077043** | Duration of the day (SD) |
| **0.09799** | Average acceleration during inactivity during the sleep period (SD) |
| **0.269356** | Sex |
| **0.392258** | (intercept) |

For the full model including age and sex, 59 actimetry-derived traits were included in the final, best-performing model. Of these 25/59 (42.3%) of the variables were the standard deviations of the traits rather than the mean of the traits, such as the time spent sedentary, sleep onset time, time walking, and the total sleep time. For 11 traits, both the means and the variances of these traits were predictive of ASD status: average acceleration during inactivity during wake, duration of light activity in bouts of 10 minutes, duration of unbouted light activity, duration of accumulated inactivity blocks during the day, M10 average peak light, time in mixed physical activity, number of blocks of inactivity, number of blocks of light activity, sedentary time, sleep duration at night, and vigorous physical activity. To further assess the importance of traits means to their variance, we generated separate elastic net models for mean values and corresponding standard deviations. There was no significant difference between the AUCs between means only and variance only evaluations (Fig. 1e), confirming that the variance of the traits may be of roughly similar importance in predicting ASD status as the means of the traits. The majority of the selected traits from GGIR and Accelerometer associated with ASD were higher variance or mean of inactivity and lower physical activity (light, moderate, and vigorous) measures.

### Relationship between Sleep and Physical Activity in ASD

Sleep and physical activity have previously been shown to correlate, with higher physical activity associating with higher reported sleep quality ^22,61^. Here, we evaluated how ASD status may affect the relationships between physical activity and sleep. First, rank correlations were performed to correlate physical activity traits and sleep traits (Fig. 2a). Overall, individuals with higher average total inactivity time or higher time spent sedentary had lower duration of sleep and a later onset of sleep (Fig. 2a).

**Figure 2.**
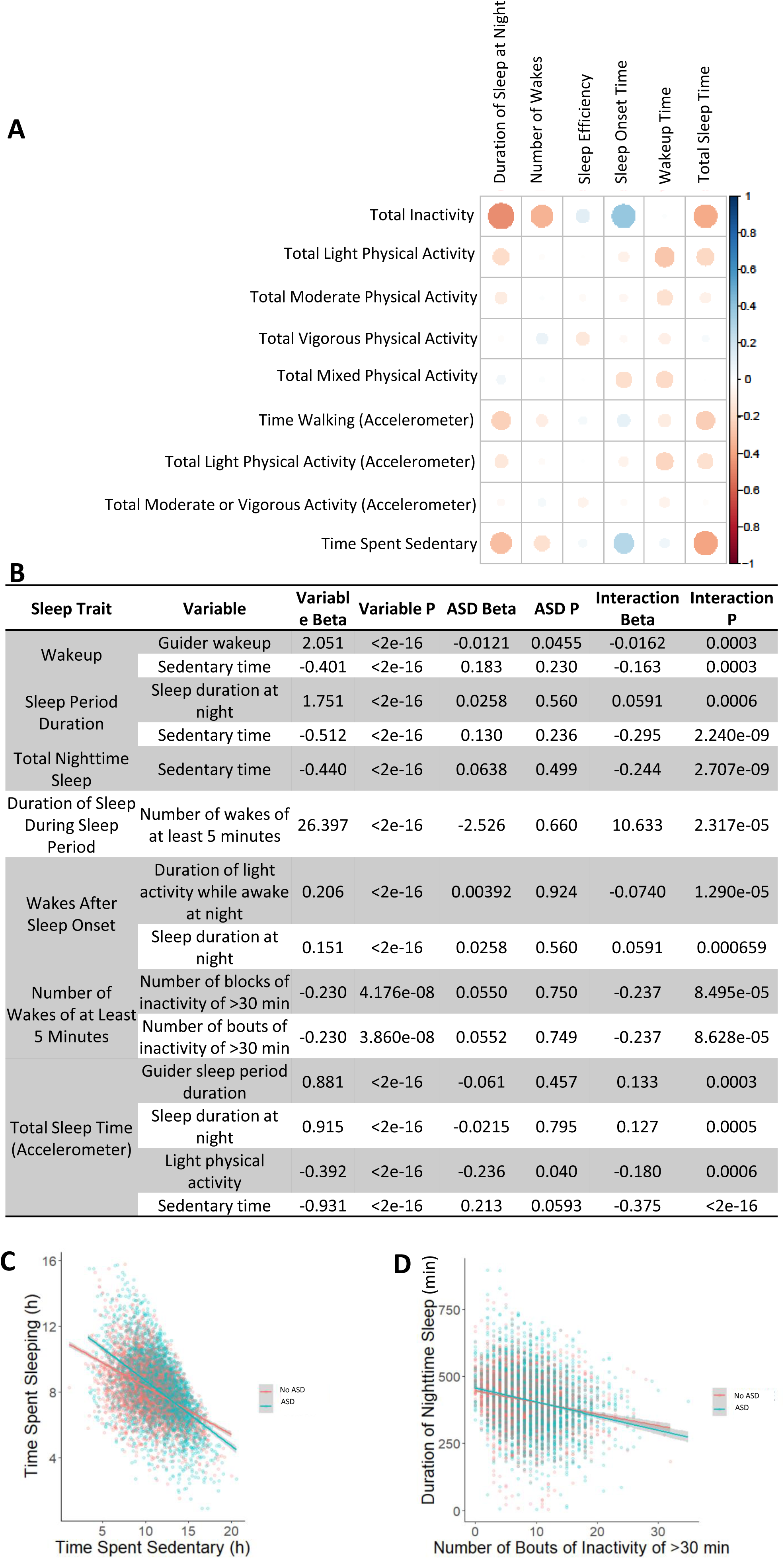
Relationship between physical activity and sleep in ASD. A) Rank correlation matrix for average physical activity traits compared to average sleep traits. Total inactivity and sedentary behavior correlated negatively with duration of sleep and positively with time of sleep onset. B) Table of significant interactions between elastic net-selected actimetry-derived traits with ASD, and sleep traits. Linear mixed effects regressions were performed with all nights for all ASPE individuals for elastic net-selected actimetry-derived traits with controls for age, age^2^, sex, and individual/family differences. Each variable significantly predicted sleep time, but also interacted with ASD status in predicting sleep time. C) The relationship between total sleep time and time spent sedentary, split by ASD status. D) Interaction of number of bouts of inactivity of at least 30 minutes and total nighttime sleep with ASD.

To evaluate how ASD may modulate this relationship, we utilized linear mixed effects regressions with interaction terms for all days (n = 6,878) for all ASPE individuals (n = 192 ASD and 223 no ASD) that passed quality control in the machine learning. Briefly, we analyzed elastic-net determined actimetry-derived traits of importance to the sleep traits, relative to ASD status. For covariates to control for confounding, we included age, sex, and age^2^ as fixed effects, and two random effects variables for individual and family effects.

We used all ten sleep traits as the outcome variables in linear mixed effects regressions. To limit the number of evaluated predictor variables, we selected only the actimetry-derived traits that were included in the best-performing machine learning model (n = 48; excluding overlapping traits that had both means and SDs). As we used all nights, rather than the aggregated nights, trait scores were not the mean or the standard deviation; they were the score for that day or night. Briefly, seven of the ten sleep traits had at least one significant interaction between an actimetry-derived physical activity trait and ASD status (Fig. 2b; Supplementary Table IV). For example, total sleep time (from the Accelerometer algorithm) showed an interaction with the time spent sedentary and ASD status, where ASD individuals had a larger reduction in total sleep time when they have a longer period spent sedentary than their non-ASD counterparts (Fig. 2b,c). Specifically, for every 1 hour of sedentary time, individuals with ASD had on average 24.84 minutes less total sleep time, whereas for every 1 hour of sedentary time, individuals without ASD had only 17.70 minutes less total sleep time. After controlling for age, age^2^, sex, individual and family differences, ASD individuals had an average decrease of 29.71 minutes of sleep for each hour of sedentary time, while those without ASD had a decrease of 20.85 minutes on average. Taking into account the mean time spent sedentary for our sample (10.978 hours), ASD individuals may have between 70-100 minutes of sleep less than their non-ASD relatives for remaining sedentary on average. As accelerometer differentiates between sedentary behavior and sleep, this cannot be explained by daytime naps. Similarly, bouts of inactivity of at least 30 minutes interacted with ASD when predicting nighttime sleep (Fig. 2b,d).

### Evaluation of Actimetry-Derived Traits Association with ASD-related Phenotypes

For all ASPE subjects, we also collected ASD-linked phenotype data using the Broad Autism Phenotype Questionnaire (BAPQ) ^50^, the Social Responsiveness Scale (SRS) ^48^, and the Behavior Rating Inventory of Executive Function (BRIEF) through informant report ^51^. A correlation analysis was conducted between the BRIEF, SRS, and BAPQ data (Fig. 3a&b).

**Figure 3.**
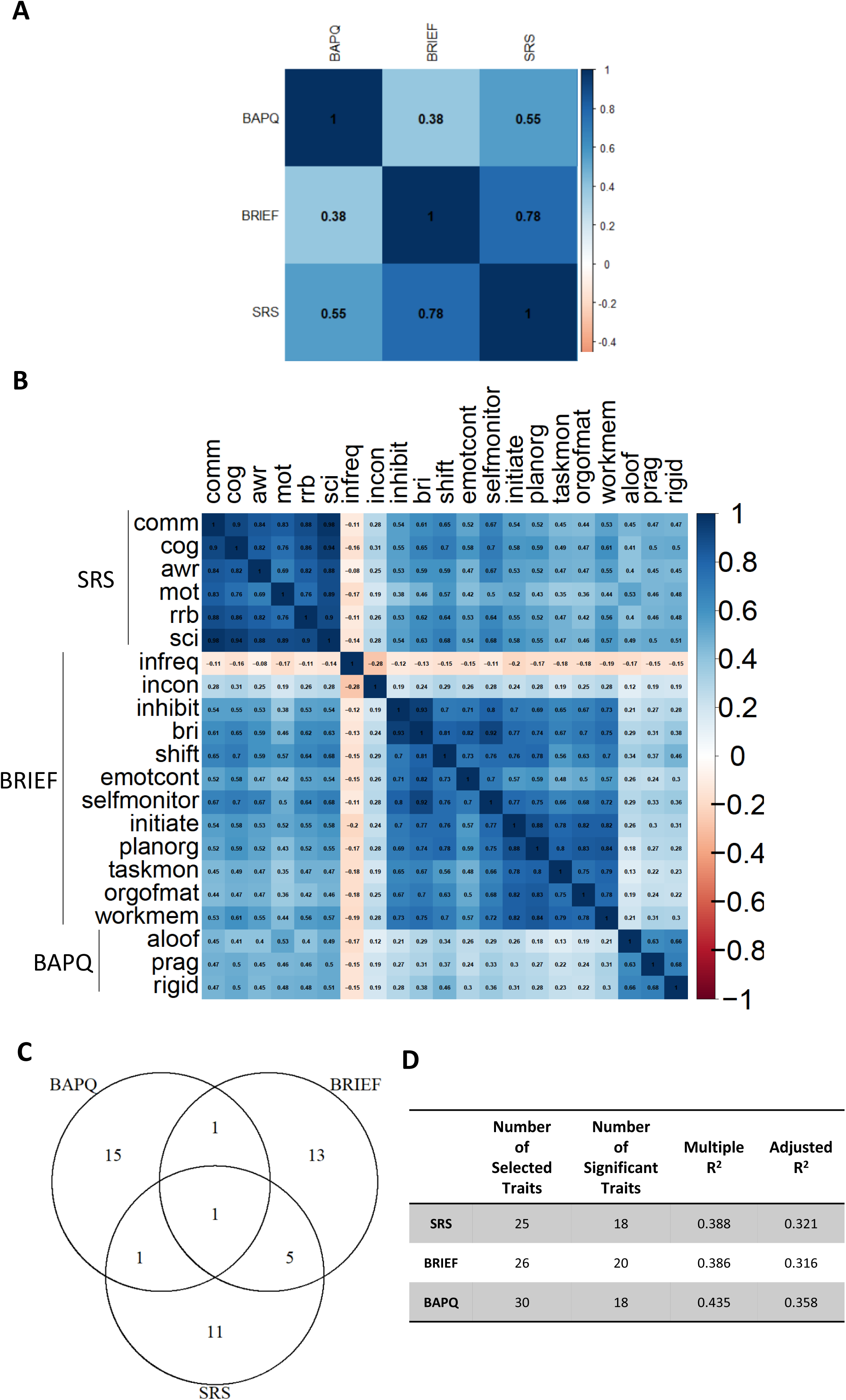
Phenotypic correlation of ASD and accelerometer-derived traits: A) A correlation matrix showing the correlations between ASD traits assessed based on three questionnaires: the Broad Autism Phenotype Questionnaire (BAPQ), the Behavior Rating Inventory of Executive Function (BRIEF), and the Social Responsiveness Scale (SRS). B) The correlation matrix for the subscales of these ASD trait questionnaires. C) The table shows the number of actimetry-derived sleep and activity traits that were significant after stepwise regression for prediction of ASD status with a model performance shown as R^2^. D) Venn diagram showing the overlap of significant actimetry-derived traits between the ASD-related assessments.

We also used a bi-directional, stepwise regression to identify significant actimetry-derived traits that were predictive of BAPQ, SRS, and BRIEF scores in all participants, including those family members without ASD, similarly to our previous analyses ^44,45^. Out of the 214 total accelerometer-derived rest/activity characteristics, 18, 20, and 18 significantly predicted ASD status for SRS, BRIEF, and BAPQ, respectively (Fig. 3c). These models explained 32%, 32%, and 36% of the variance within SRS, BRIEF, and BAPQ, respectively (Fig. 3c). When investigating the trait overlap, one trait was selected for all three questionnaires: the standard deviation between days of the average amount of acceleration during inactivity (Fig. 3d), where a higher variance in inactivity was associated with higher SRS, BRIEF, and BAPQ scores. SRS and BRIEF had the most overlap in selected traits, with 6 actimetry-derived traits common between them: the variances of the unbouted activity during the day and inactivity while awake during the sleep period, the total moderate physical activity while awake during the sleep period, the variance of total moderate or vigorous physical activity, the number of bouts of moderate or vigorous activity between 5 and 10 minutes, and the average acceleration during moderate or vigorous activity between 5 and 10 minutes (Supplementary Table V).

### Validations of findings in the SPARK collection

Using machine learning, we showed the importance of physical activity traits and their relationship to ASD status in the ASPE cohort. To validate these findings and further investigate the relationship between sleep traits and ASD, we increased the number of ASD subjects by including the subjects from the SPARK collection. We used the same 3-week protocol for data collection and quality control that was used for the ASPE participants. Two representative actimetry double-plots, plotted using the ChronoSapiens package^62^ and visualizing overall activity during the three weeks of actimetry, show a highly regular and highly irregular activity pattern (Fig. 4a,b). SPARK participants were further separated based on the presence or absence of sleep problems, as determined by the SPARK medical questionnaire. Overall, SPARK individuals with ASD were much more likely to report sleep problems than family members without ASD (Fig. 4c,d). Next, participants were separated into these groups: a) ASPE ASD probands (AP); b) ASPE relatives without ASD and unaffected by other psychiatric conditions (UR), c) SPARK ASD probands with reported sleep problems (SP (wSP)), d) SPARK ASD probands without reported sleep problems (SP (noSP)). Overall, the UR and AR groups were older than the other groups (Fig 5a). To evaluate how reported sleep problems may associate with sleep regularity, we employed the sleep regularity index (SRI)^63^. Briefly, a higher score suggests more regularity in their sleep problems, while a lower score suggests more irregular sleep. Individuals with ASD, both in ASPE and SPARK, had significantly lower SRI scores than the ASPE relatives with no psychiatric conditions (Fig. 4e). Additionally, those SPARK individuals who reported sleep problems had an even lower regularity in their sleep values (Fig. 4e). Correlations between the SRI and physical activity were performed showing that higher light activity, lower inactivity, and higher moderate physical activity correlated to more regularity in sleep, with individuals with ASD both from SPARK or ASPE having overall lower SRI at the same level of physical activity for inactivity and light activity (Fig. f,g,h).

**Figure 4.**
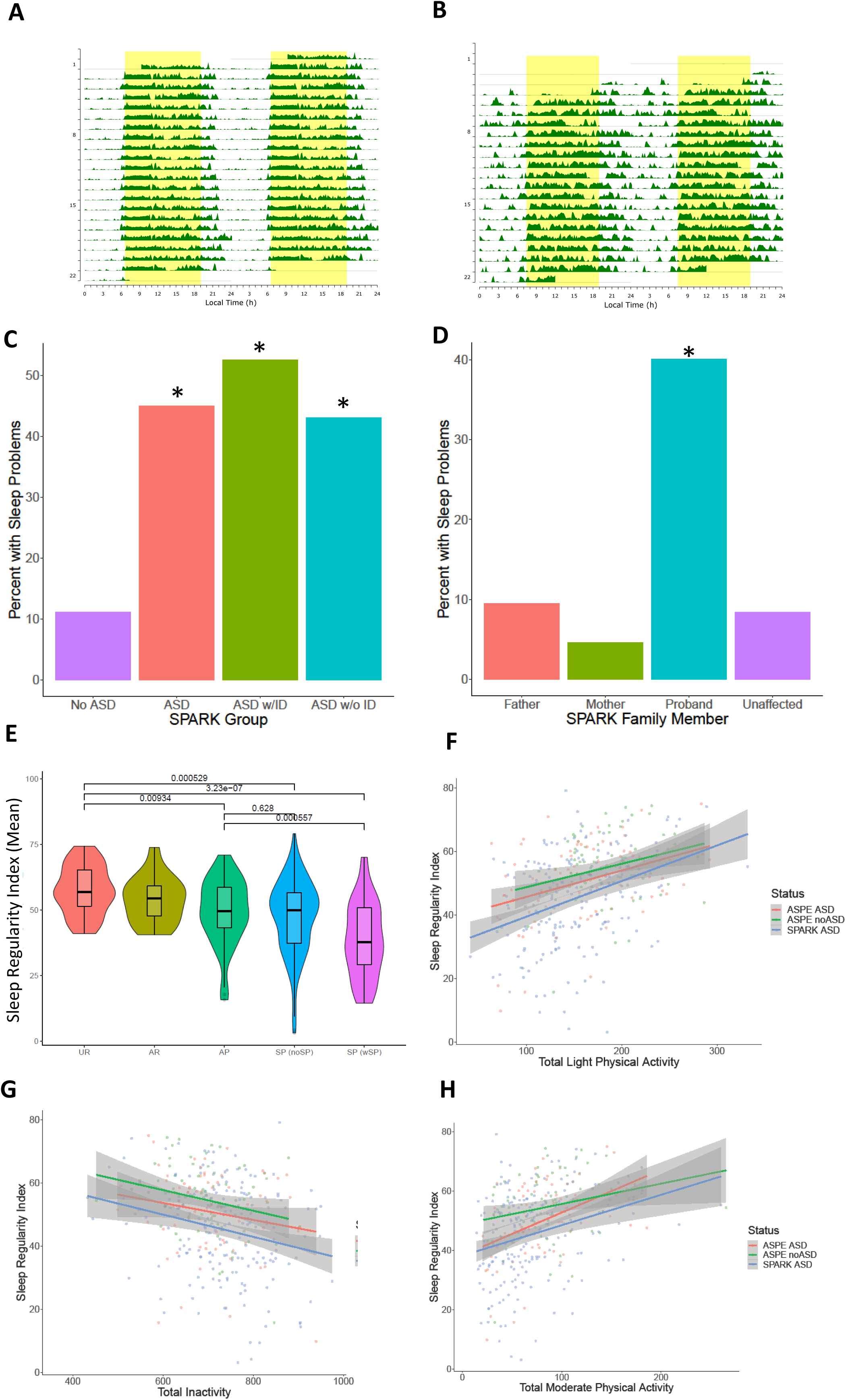
Sleep regularity and reported sleep problems in the SPARK cohort. Actimetry-derived double plots visualized using the ChronoSapiens^62^ package. These plots show the activity throughout the three weeks of data collection, where A) shows a highly regular activity pattern individual and B) shows a highly irregular activity pattern individual. C) Reported sleep problems in the SPARK medical questionnaire. Individuals with ASD report higher levels of sleep problems than their non-ASD family members (p < 0.005). D) In European ancestry quartets, where data is available for both parents and an unaffected sibling, the ASD probands are more likely to report sleep problems than any other family member role. E) Of our recruited SPARK participants, 25.6% reported sleep problems while 74.7% did not in the SPARK questionnaire. F) When comparing the sleep regularity index (SRI) scores between groups, the individuals with ASD were significantly more irregular in their sleep than the ASPE unaffected relatives. The SPARK individuals with reported sleep problems were also more irregular than the ASPE ASD individuals. UR: ASPE unaffected relatives, AR: ASPE relatives with a psychiatric diagnosis other than ASD, AP: ASPE ASD probands, SP (noSP): SPARK probands with ASD with no reported sleep problems, SP (wSP): SPARK probands with ASD with reported sleep problems.

**Figure 5.**
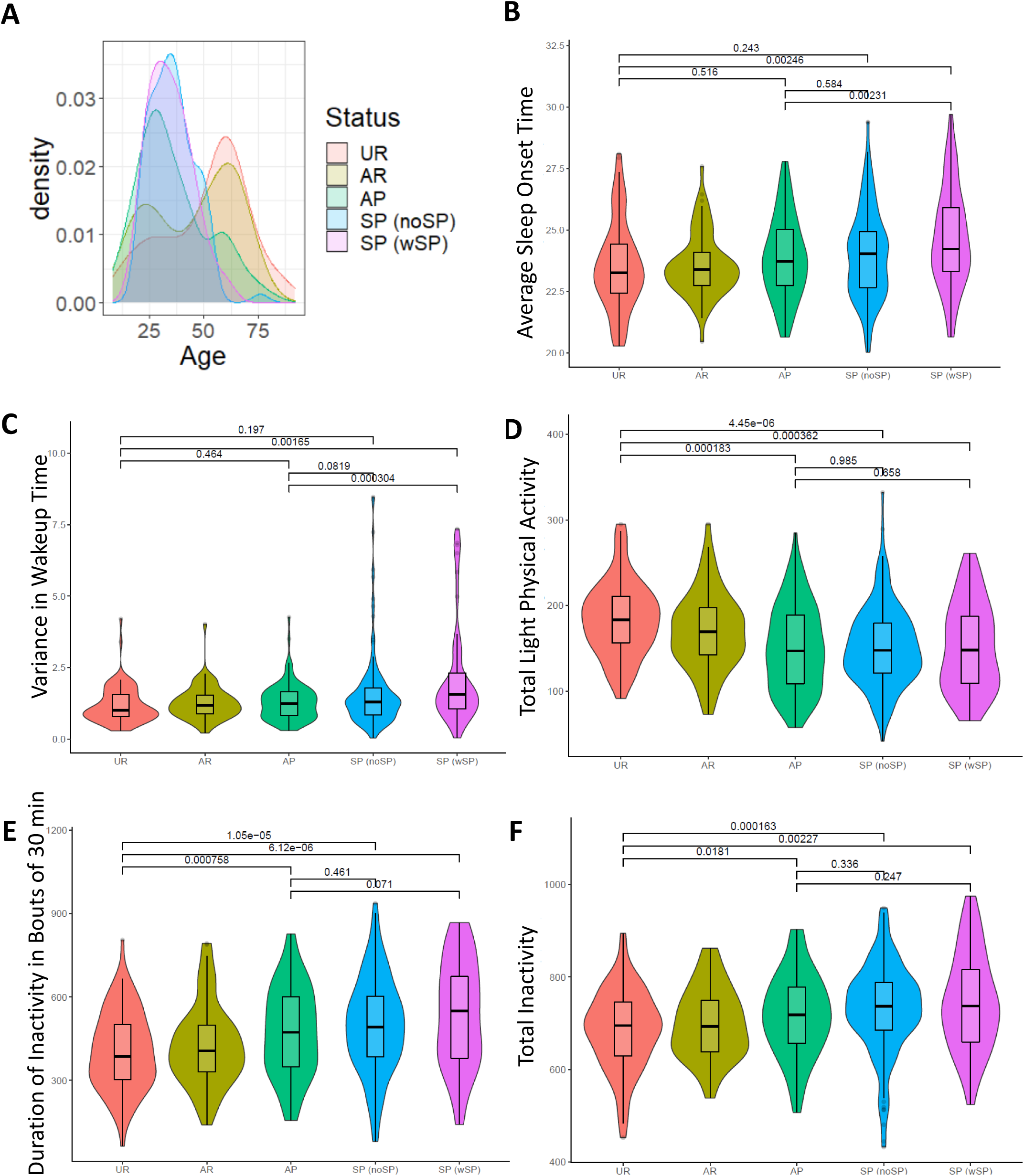
Sleep and physical activity comparisons in ASPE and SPARK participants. A) Age distributions of ASPE unaffected relatives (UR), ASPE relatives with a psychiatric diagnosis other than ASD (AR), ASPE ASD probands (AP), and SPARK probands with ASD with (SP (wSP)) or without (SP (noSP)) reported sleep problems. Violin plot comparisons between the three groups, with p-values corrected for age, age^2^, and sex. B) Average sleep onset between the groups, C) Variance in wakeup times, D) total light physical activity, E) duration of inactivity in bouts of 30 minutes or more, F) total inactivity.

As above, we compared sleep and activity measures in the AP, UR, SP (wSP), and SP (noSP) groups. A comparison of all 218 actimetry-derived GGIR traits using linear regressions controlling for age, age^2^, and sex showed that SPARK and ASPE ASD subjects were significantly different from the non-ASD ASPE relatives in several physical activity traits, including total light activity and long periods of inactivity and the intra-individual variance of wakeup time, where individuals with ASD had higher inactivity times and lower light activity than the ASPE family members without ASD (Fig. 5b-f).

## DISCUSSION

Our results indicate that the difference in several sleep and motor activity traits (the means or the variances) can differentiate individuals with ASD from those without ASD. These findings emphasize the importance of not only looking at the averages of traits, but the variances as well in longitudinal studies. Additionally, our findings stress the importance of not only investigating sleep, but also physical activity in ASD. Our accelerometer-based study of 382 ASD subjects (ASPE and SPARK) and 223 non-ASD relatives (ASPE) using three weeks of continuous measures revealed that physical activity traits, including day-to-day variability of these traits, are predictive of ASD status in this family-based sample. What sets our study apart is the substantial representation of adult subjects with ASD, in contrast to many previous reports focusing primarily on sleep and physical activity traits in children with ASD. Also, to reduce heterogeneity and influence of repetitive movements, our study primarily focuses on adults who do not present with intellectual disability (ID). This study design also permitted the comparison of self-reported and device-derived sleep assessments.

In this study, we used machine learning methods to evaluate 242 total actimetry-derived sleep:wake characteristics, and their potential associations with ASD in a high-throughput, unbiased approach. In the best-fit model, nearly half of the selected traits were the day-to-day variability of an actimetry-derived trait rather than their mean values, suggesting the importance of examining the day-to-day variability as well as general average differences in sleep and physical activity in ASD. The relationship between these selected traits and several measures of sleep quality were also evaluated for potential interaction with ASD status. Here, we found that total time spent sedentary had a different relationship with total sleep time in individuals with ASD compared to those without ASD, where individuals with ASD had more sleep loss for the same amount of sedentary time than their non-ASD relatives. This may suggest a different relationship between physical activity and sleep in ASD subjects, where physical activity is more strongly tied to sleep in those with ASD, a finding with potential treatment implications. Future research to parse out the relationship between physical activity, sleep, and ASD is warranted.

Interestingly, many of the traits selected from the elastic net were physical activity rather than sleep, further supporting the importance of general level or pattern of movement in ASD. Other researchers have found that deficits in motor skills in children and adolescents with ASD can predict social deficits ^64,65^. The relationship between these selected traits and several measures of sleep quality were also evaluated, with the potential interaction with ASD status. Here, we found that total inactivity and total light activity during the day had a stronger correlation with total sleep time in individuals with ASD than their unaffected relatives As further support for the relationship between sleep and physical activity relationship being important in ASD, we observed that individuals with ASD at overall lower sleep regularity, but in those individuals (and in individuals without ASD), higher light and moderate physical activity and lower inactivity associated with more sleep regularity.

Day-to-day intra-individual variability in sleep and physical activity phenotypes appeared to be predictive of ASD status in our samples. Other researchers have also found that variability within sleep efficiency and number of awakenings is significantly correlated with ASD symptoms ^66^, further suggesting the importance of the collection of longitudinal, multi-day datasets to evaluate the variability of traits and the relationship of these phenotypes with the core ASD symptoms. Studies of sleep in both *Drosophila melanogaster* and in Genetic Reference Panel and Diversity Outbred mice demonstrated that, at least in part, genotype influences day-to-day variability in sleep characteristics ^67,68^. The importance of these findings is also supported by studies showing that a high degree of day-to-day variability may represent a hallmark of both sleep disorders and overall mental health, associated with attention deficit disorder (ADHD), post-traumatic stress disorder (PTSD), and changes in mood or level of depression ^69–73^. A prior study also found that algorithms could distinguish between ADHD and non-ADHD subjects using intra-individual day-to-day variability, rather than the mean differences in traits ^74^. More broadly, a recently published consensus report by the National Sleep Foundation found that consistency of sleep onset and offset timing is important for health, safety, and performance^75^.

Through our study, we have provided further evidence of the importance of physical activity measures in ASD. This agrees with previous findings suggesting that physical activity interventions may decrease symptoms of ASD ^26^. It is, however, difficult to parse out whether this is a direct effect of the physical activity or if the physical activity itself may improve sleep function, thereby improving ASD symptoms. The interplay between physical activity and sleep is likely bidirectional^22^.

Our study has several limitations. First, in addition to the need to extend our study to a larger set of ASD individuals, we also need an independent set of unrelated non-ASD subjects to confirm our findings. These data would also strengthen the machine learning approach by providing further controls against model overfitting. To evaluate both sleep and physical activity in ASD subjects, we opted to perform an accelerometer-based study combined with self-reports. However, polysomnography (PSG) is considered the gold-standard assessment for sleep/wake disturbances and electrophysiological evaluation of sleep stages. For example, multiple lines of evidence suggest that sleep spindles, a characteristic electroencephalogram (EEG) signature of stage 2 non-rapid eye sleep, may represent heritable biomarkers of a range of neuropsychiatric disorders and may be influenced by dynamic alterations in synaptic plasticity seen during sleep ^76^. Ideally, future large-scale investigations of sleep architecture in ASD will involve a range of sensors including non-invasive EEG monitoring ^77,78^. Also, simultaneous monitoring of heart rate variability, which has been shown to associate with psychiatric conditions, may provide insights into the ASD pathophysiology ^79^.

Continued research into the sleep and activity patterns of autistic individuals should include the evaluation of day-to-day variability and physical activity measures, as these appear to be strong indicators of ASD status. Previously, researchers have investigated the relationship between physical activity and sleep, but our findings of an interaction in this relationship with ASD status support the utility of more research in the autism space. Taken with prior reports showing improvement of ASD symptoms after interventions designed to improve physical activity, long-term studies evaluating the relationships between sleep, physical activity, and ASD should be conducted to evaluate the potential therapeutic benefits of intervention in these domains.

## Author Contributions

JDW was involved in the formulation of the project, analysis of actimetry data, ran statistical analyses, wrote, and edited the manuscript. AG was involved in the collection and analysis of actimetry data and the formulation of the project. MF was involved in the formulation of the project, edited the manuscript, and helped lead the research team. BG, MR, and HCD recruited participants, administered ASD-related trait questionnaires, and distributed actimetry watches. DJR, LA, ESB, RS, and MB formulated the project, edited the manuscript, and led the research teams. BTK assisted with statistical analyses and edited the manuscript.

## Acknowledgments

We would like to thank all participants and investigators in Autism Spectrum Program of Excellence (ASPE) and Simons Powering Autism Research (SPARK). We would like to thank the members of the ASPE and SPARK research teams for their assistance. We also thank Matthew Kayser, Ilan Dinstein, Till Roenneberg, and Jonathan Mitchell for their advice, and Lis Zandbergen for her assistance in analyzing the data.

This work was supported by NIH grant (NIMH R21 MH093415 to MB), Simons Foundation Award (ID: 877185) as well as the Autism Spectrum Program of Excellence (ASPE) at the University of Pennsylvania.

