## SupplementaryIndex for "Sleep and Activity Patterns in Autism Spectrum Disorder"

### Supplementary Tables and Figures

- Supplementary Table I: Demographics of ASPE and SPARK participants. (see below)
- Supplementary Table II: List of all actimetry-derived traits from GGIR and Accelerometer. (SupTable2_Actimetry_Traits.csv)
- Supplementary Figure 1: Heatmap of correlations between actimetry-derived traits. (see below)
- Supplementary Figure 2: Evaluation of the alpha hyperparameter (see below)
- Supplementary Figure 3: Comparison of seasonality of data between ASD and non-ASD groups. (see below)
- Supplementary Figure 4: Comparison of weekdays and weekend days in even/odd datasets, and in ASD vs. nonASD groups. (see below)
- Supplementary Table III: Number of nights for each season. (see below)
- Supplementary Table IV: Elastic net selected actimetry-derived traits without age and sex. (SupTable4_elasticnettraits_noagesex.csv)
- Supplementary Table V: Linear mixed-effects regression results for all ten sleep traits, using elastic net-selected actimetry-derived traits. (SupTable5_LMER.xlsx)
- Supplementary Table VI: ASD-related questionnaire stepwise regression results (SupTable6_AllActVariables.csv)
- Supplementary Table VII: All GGIR trait comparisons with ASPE and SPARK (SupTable7_ASPE_SPARK_Comparisons.csv)

### Supplementary Table I

| | ASPE ASD (n = 190) | ASPE Family Members without ASD (n = 223) | SPARK ASD Probands (n = 190) |
|---|---|---|---|
| Age, mean (sd) | 35.7 (17.2) | 45.6 (19.5) | 35.0 (9.86) |
| Sex, female (%) | 74 (38.5%) | 127 (57.0%) | 54 (28.4%) |
| BAPQ, mean (sd) | 4.16 (0.76) | 2.6 (0.69) | - |
| SRS, mean (sd) | 0.66 (0.84) | -0.52 (0.72) | - |
| BRIEF, mean (sd) | 0.521 (0.85) | -0.43 (0.84) | - |

Supplementary Table I. Demographics information for all participants

Supplementary Figure 1

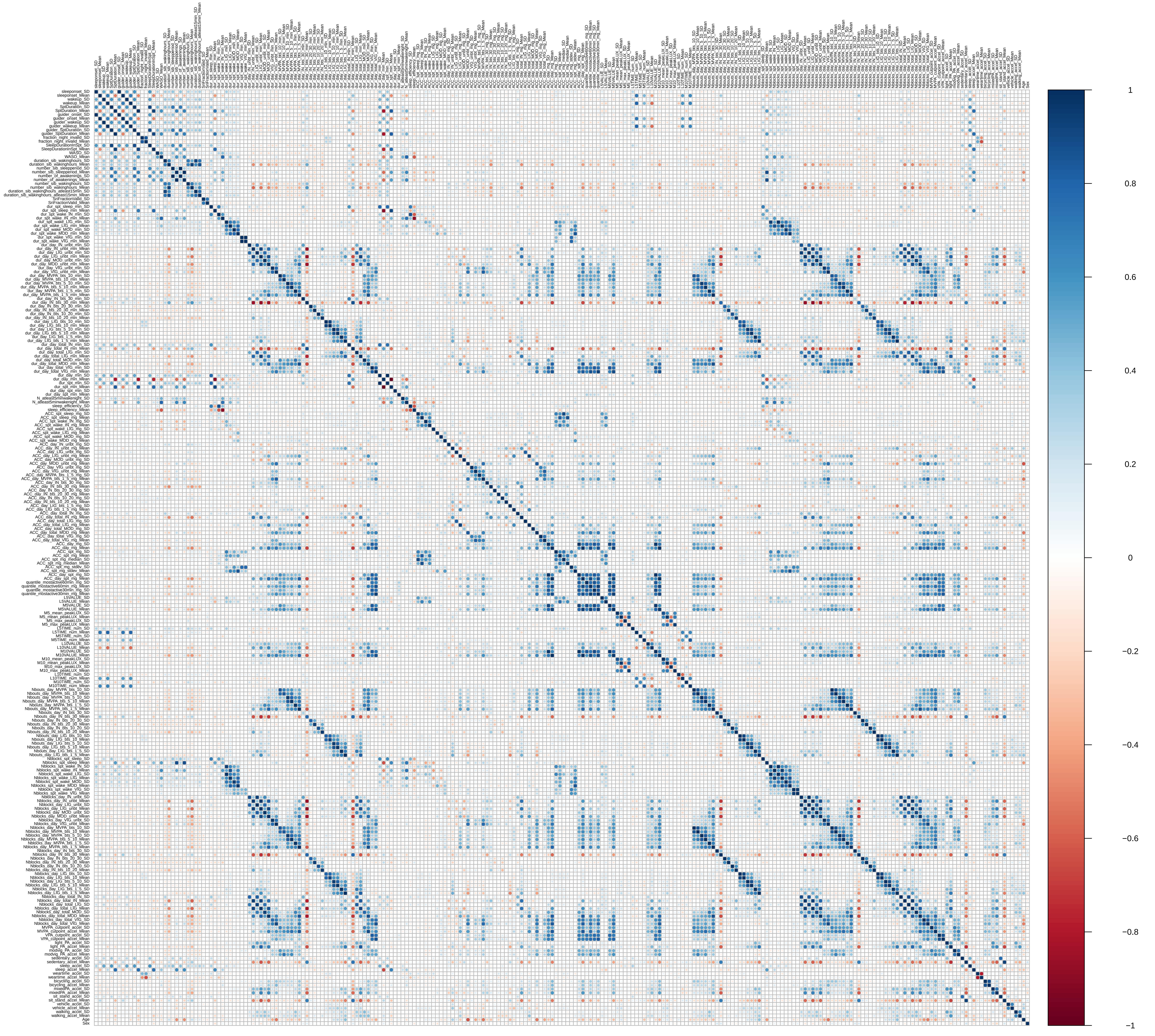

### Supplementary Figure 2

**A**

**B**

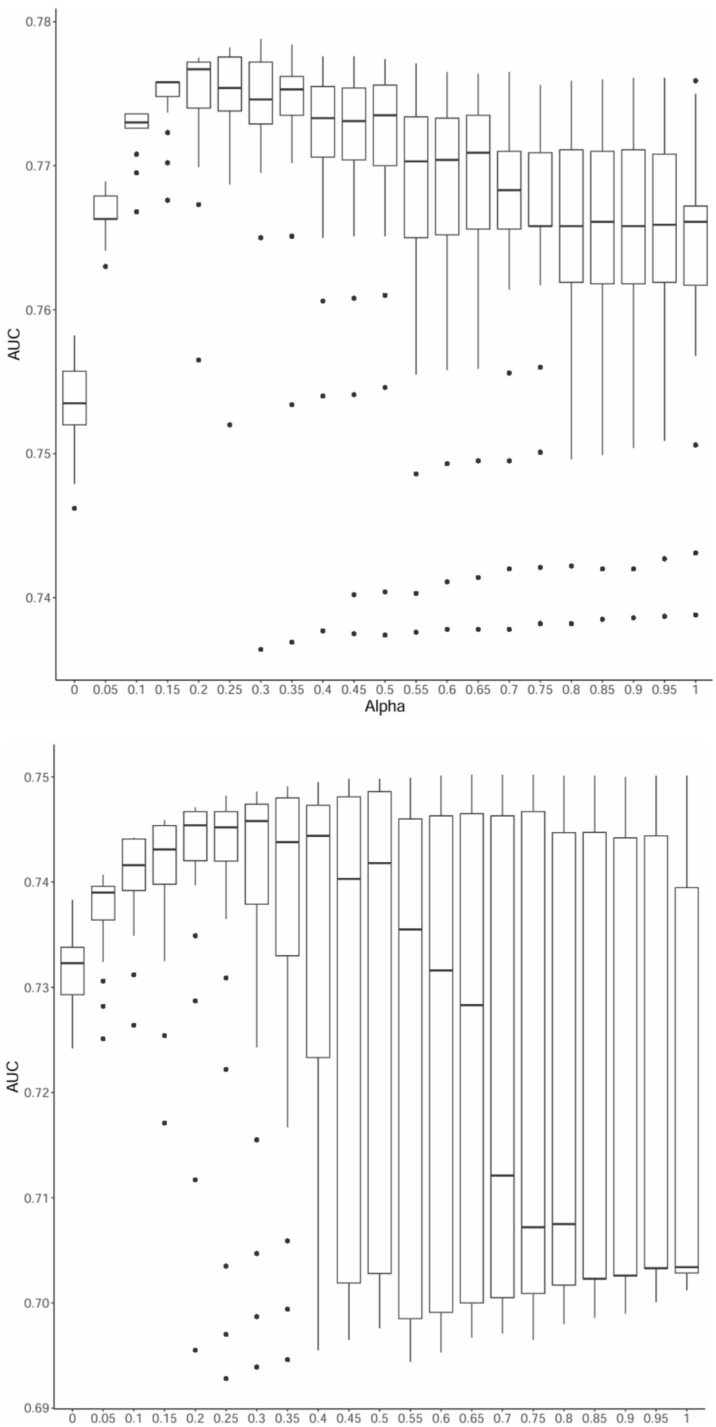


Supplementary Figure 2. Elucidating the best alpha value for machine learning. In A) with age and sex in the model, B) no age and sex in the model.

### Supplementary Figure 3

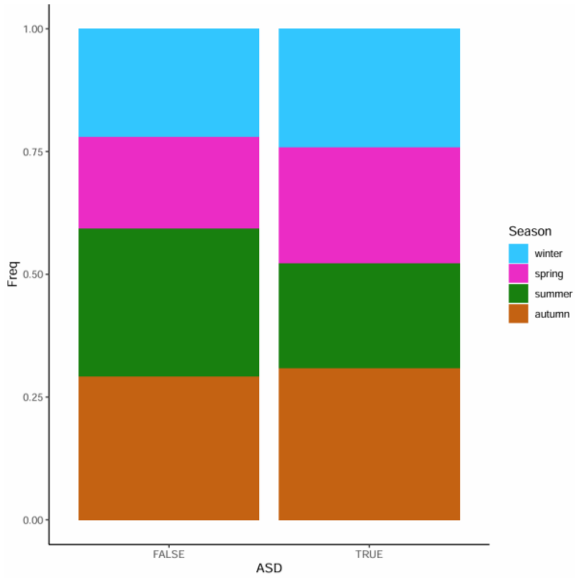


Supplementary Figure 3: Season of actimetry recording in ASD and non-ASD individuals. Individuals without ASD were more likely to be assessed with actimetry in summer than those with ASD. ASD individuals were more likely to have actimetry measures in the springtime.

### Supplementary Figure 4

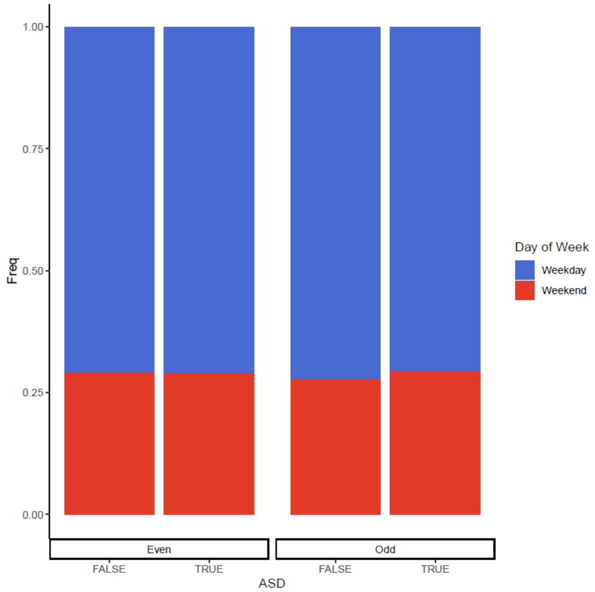


Supplementary Figure 4: Distribution of weekdays and weekends in those with and those without ASD. There was no significant difference between ASD and non ASD individuals for weekdays and weekends, even after splitting even and odd days. No difference in distribution in even or odd datasets ($p > 0.5$).

#### Supplementary Table III

| | Spring | Summer | Autumn | Winter |
|---|---|---|---|---|
| **ASD** | 536 | 480 | 698 | 543 |
| **No ASD** | 704 | 1,122 | 1,093 | 821 |

Supplementary Table III. Number of nights within each season, for ASD and non ASD participants. ASD participants were more enriched for winter over summer ($p = 2.4e-8$)
